# Synthqa - Hierarchical Machine Learning-Based Protein Quality Assessment

**DOI:** 10.1101/2021.01.28.428710

**Authors:** Mikhail Korovnik, Kyle Hippe, Jie Hou, Dong Si, Kiyomi Kishaba, Renzhi Cao

## Abstract

**Motivation:** It has been a challenge for biologists to determine 3D shapes of proteins from a linear chain of amino acids and understand how proteins carry out life’s tasks. Experimental techniques, such as X-ray crystallography or Nuclear Magnetic Resonance, are time-consuming. This highlights the importance of computational methods for protein structure predictions. In the field of protein structure prediction, ranking the predicted protein decoys and selecting the one closest to the native structure is known as protein model quality assessment (QA), or accuracy estimation problem. Traditional QA methods don’t consider different types of features from the protein decoy, lack various features for training machine learning models, and don’t consider the relationship between features. In this research, we used multi-scale features from energy score to topology of the protein structure, and proposed a hierarchical architecture for training machine learning models to tackle the QA problem.

**Results:** We introduce a new single-model QA method that incorporates multi-scale features from protein structures, utilizes the hierarchical architecture of training machine learning models, and predicts the quality of any protein decoy. Based on our experiment, the new hierarchical architecture is more accurate compared to traditional machine learning-based methods. It also considers the relationship between features and generates additional features so machine learning models can be trained more accurately. We trained our new tool, SynthQA, on the CASP dataset (CASP10 to CASP12), and validated our method on 33 targets from the latest CASP 14 dataset. The result shows that our method is comparable to other state-of-the-art single-model QA methods, and consistently outperforms each of the 14 used features.

**Availability:** https://github.com/Cao-Labs/SynthQA.git

## 1 Introduction

The accurate prediction of the true structure of a protein from protein sequences has long been a focus of research in the Bioinformatics field [1, 2, 3, 4, 5, 6, 7, 8, 9]. This is because the ability to accurately predict these microscopic structures would enable many valuable improvements in the pharmaceutical drug and vaccine design process and advancements in the design and creation of new proteins for specific functions [10, 11]. Unfortunately, discovering the true structures of proteins is not easy. Most of the protein structures are determined by experimental techniques, such as X-ray crystallography and Nuclear Magnetic Resonance, but those techniques are extremely time-consuming, expensive, and sometimes even impossible [12]. Because of this, the total number of proteins with native structure is still few and far between compared with the vast number of existing proteins. In response, research in recent years has turned its focus to the application of machine learning techniques to assist with this problem and aid in the reliable and timely prediction of tertiary protein structures.

Computational protein structure prediction methods with the help of machine learning techniques usually work on two main problems: model sampling and model ranking [10]. Model sampling is the process by which protein structural models are generated using machine learning prediction algorithms or other template-based methods searching against the database. On the other hand, model ranking is concerned with quality assessment (QA) of protein structural models generated via model sampling. The goal of QA is to rank these models and select the best ones to be the final predictions. To do this, we assume the native structure is not yet available, and models are analyze and compare the models with the protein target’s true native structure to help the AI learn the pattern [10]. The model ranking problem is what we focus on in this research.

There are two strategies to solve the model ranking problem. The consensus (also known as clustering) approach involves the QA system considering many generated models from a variety of prediction algorithms and searching out patterns within the models to predict which model is most accurate. The consensus model then outputs the relative quality scores for each model using pairwise structural comparison [13]. In contrast, the single-model approach focuses on one generated protein model at a time, making it time-optimal for processing large data sets. Researchers have already conducted many studies and experiments to tackle this protein prediction QA problem (e.g, the CASP experiment is designed to validate the accuracy of computational protein structure prediction methods every other year since 1994 [14]). For example, one consensus-model QA experiment called MULTICOM [10] was a particularly successful method, combining an unprecedented 14 different QA methods into one diversified system. Its performance won third place out of 143 contestants in the CASP11 contest, a notable achievement.

On the other hand, past single-model QA experiments have tried integrating traditional machine learning techniques, such as DeepQA [15], which used a Deep Belief Network to process 16 different features of proteins such as energy, physiochemical characteristics, and structural information. DeepQA outperformed support vector machines, neural networks, and ProQ2 [16], with a notable strength in ranking ab initio models. Less conventional methods like Qprob [17] focused on finding the absolute error of each features’ value and comparing those against the true score (GDT-TS or GDT score) of the corresponding structural model, ultimately estimating the probability density function to make its predictions. Like DeepQA, Qprob did well in the CASP experiment for single-model QA methods. Other methods, such as the unique SMOQ [18], evaluated absolute residue-specific local qualities of a single protein model. Unlike other QA programs, SMOQ converted its predicted local qualities into a single score to predict a model’s global quality, making it unique in both methodology and output. The result was comparable to many top single-model QA methods. The SVMQA [19] method is also a notable single-model method consisting of a support vector machine trained on statistical potential energy terms and consistency-based terms between structural features. It calculated the TM-score and GDT scores, and was especially adept at selecting high-quality models from decoy data. Moreover, the new method VoroMQA [20] uses statistical potentials combined with interatomic contact areas to perform quality assessment of protein models which makes it unique. It outputted both local and global quality scores for any protein decoy and performed very well in the recent CASP experiment.

On a general level, the consensus approach has shown better performance than single-model methods in past CASP experiments, especially on large datasets [21]. However, consensus methods depend a great deal on the size and quality distribution of their input models, making it difficult to find the best models in most cases, especially if the best model does not happen to be the average one and bears much similarity to the other models [13]. In other words, if there are several models within the pool that are not like the others then the consensus methods have a hard time distinguishing between the like models and the outliers. This makes it ever more important to develop single-model prediction methods that are not reliant on any other models to predict a model’s quality.

Deep learning has become a fascinating area of study when it comes to protein model quality assessment problems. DeepQA [15] and other deep learning methods [22, 4, 14] like ProQ3D [23], DeepTracer [24], 3DCNN [25] have a tendency to outperform other traditional machine learning methods (e.g., support vector machines, neural networks, etc.) in the Bioinformatics field, making deep learning as a QA strategy an important avenue of study to pursue. As mentioned above, DeepQA uses a Deep Belief Network and utilizes the weights of two Restricted Boltzmann machines in its network which are adjusted throughout the training process. ProQ3D uses the same carefully selected and optimized input features as ProQ3, but replaces ProQ3’s support vector machine with a deep neural network which ultimately achieved very good performance. 3DCNN attempts to learn the features of a predicted protein structure directly from the raw data of the CASP7 to CASP10 datasets and it ranked the protein structure decoys accurately using a deep convolutional network based on the raw three-dimensional atomic densities. However, most of the protein model quality assessment methods didn’t consider different type of features or the relationship among those input features. The deep learning technique may learn those relationships, but it might be difficult and could consume a lot of computational resources. In this research, we consider multi-scale features for the protein model quality assessment problem and propose a hierarchical architecture for training machine learning models which considered the relationship of features and trained the machine learning model layer by layer with a much easier way to understand.

Description of features selected for training along with their assigned labels.

|  |  |  |
| --- | --- | --- |
| Feature Label | Value Range | Description |
| EC | 0.0 - 1.0 | euclidean compact score. |
| ES | 0.0 - 1.0 | exposed surface score. |
| SU | 0.0 - 1.0 | surface score. |
| RESIDUES | 0.0 - 1.0 | residue score (extracted using SBROD). |
| HBONDS | 0.0 - 1.0 | hydrogen bonds score (extracted using SBROD). |
| SOLVATION | 0.0 - 1.0 | solvation score (extracted using SBROD). |
| BACKBONE | 0.0 - 1.0 | backbone score (extracted using SBROD). |
| ANDIS | 0.0 - 1.0 | ANDIS score. |
| R_* | 0.0 - 1.0 | rosetta energy values (6 features). |
| TQA1 - TQA512 | 0.0 - 1.0 | topology feature values. |

The paper is organized as follows. In the Methods Section, we describe the datasets and features that are used for our new machine learning method and how we process those features. In the Discussion Section, we describe the training process and performance of our new hierarchical architecture for machine learning models. We also show the performance of our method and compare it with several other QA methods. In the Conclusion Section, we conclude the paper with our findings and future works.

## 2 Methods

We considered multi-scale features and proposed a new hierarchical architecture to train a machine learning model for evaluating the quality of any predicted protein decoys. Based on previous research [15, 26, 27, 28, 17, 29, 30], our new method first generated multi-scale features which include the sequence, structural, energy and statistical potential from the input. Our new hierarchical architecture generates additional features from the initial feature sets and trains different machine learning models to improve performance. In the following sections, we describe the features that we have used in the new method and hierarchical architecture training of machine learning models for protein quality assessment problem.

### 2.1 Feature Selection

#### 2.1.1 Structural features - EC, ES, SU scores

These three scores are generated from the structure information of the decoy without any sequence input and are first introduced by researchers from Qprob and DeepQA papers ([17]; [15]). We used the same scores generated from those tools and considered that as features for the protein decoy. The details of those scores are introduced in the reference paper.

#### 2.1.2 Energy features - ROSETTA Centroid Energy Terms

Six rosetta energies described in [30] have been used recently in the research of QDeep (QDeep) for the protein model quality assessment problem. The following six energies are used as training features: residue environment (env), residue pair interactions (pair), c*β* density (cbeta), steric repulsion (vdw), packing (cenpack), and centroid hydrogen bonding (cen_hb). We chose these six energy features to understand the quality of any protein decoy, they are calculated using the tool PyRosetta4 (PyRosetta). In Table 2.1, they are referenced with the “R_” prefix (e.g. R_ENV, R_PAIR, etc.). To optimize the number of features that we use for training and for simplicity’s sake, we take the average of each energy score across all residues as the final value for each of those six features. For example, the R_ENV feature is the summation of all env values of the protein’s residues divided by the number of residues inside the protein.

#### 2.1.3 Protein Backbone Feature - SBROD

The SBROD (Smooth Backbone-Reliant Orientation-Dependent) method uses only the conformation of the protein backbone (SBROD). We extract features derived from the first stage of SBROD’s feature extraction step and run it through an arpack singular value decomposition (SVD) to reduce the size of the feature array to n by (n-1), where “n” represents the total number of input protein decoys. For example, when there are 5 protein decoys as input, we will have an array containing 5 rows and 4 columns with each row corresponding to a specific protein decoy. In the special case when only one protein decoy is provided, we duplicate the input protein decoy for generating SBROD related features to avoid a resulting array of zero columns. This leaves an array with 2 rows and 1 column to be generated for one protein decoy and both rows containing the same value. Finally, the column with the largest explained variance ratio for the decomposition is selected as the representative value for the input feature. In total, we used 4 structural interaction features from SBROD (residue, backbone atoms, hydrogen bonds, and solvation shell), the 4 feature values generated from these are used to train our SynthQA tool. The decomposition step is done to minimize the number of features and also filter the large amount of input values from SBROD so the training process of machine learning in our method is easier. The explained variance ratio is used to select the column because a column with a higher explained variance for a decomposition would suggest the values in that column best describe the original higher-dimension array, thus meaning that the column with the highest explained variance contains the most information about the protein feature than other columns in the array.

#### 2.1.4 Statistical Potential Feature - ANDIS

The knowledge-based statistical potential is an important feature for evaluating the quality of protein decoys and has been widely used for protein structure prediction. The latest research angle and distance dependent (ANDIS) statistical potential (ANDIS) considered the pairwise interaction and angles between 167 residue-specific heavy atom types. The angles are already used by researchers for the protein model quality assessment problem (e.g., AngularQA, AngularQA), and the pair-wise interaction is related to the recent research on using contact and distance information for protein structure prediction (jing2020web,hou2019protein), so that we include this statistical potential feature from ANDIS. The final score generated from ANDIS is directly used as an input feature for our method.

#### 2.1.5 Topology feature - TopQA

Although we do not use this feature in the final version of our method, we have experimented with topology features. The TopQA (TopQA) is different than any other traditional method for protein model quality assessment problem. It proposes a new representation of protein structures while the topology of the protein structure is used to evaluate the quality of this structure. We assume this would be a useful feature to represent the protein, so we extracted the 8×8×8 matrix from the TopQA tool which totals to 512 values. We convert that resulting 3D matrix to a one-dimensional array of features (each of the 512 values are in the same order as they appear in the original 8×8×8 matrix) for our new method. The contents of this array is mostly sparse with a lot of zeros. Features belonging to this output 1D array are labeled with the “TQA” prefix followed by the index (from 1 to 512) of that value in the 1D array, which is shown in Table 2.1.

#### 2.1.6 One-Dimensional Data Format

A lot of methods use two or three-dimensional input data (arrays nested inside each other) for model training. Processing such data can be complex, time-consuming, and use up large amounts of storage space. In this research, we chose to store our input training features as a single one-dimensional (1-D) array per protein decoy.

#### 2.1.7 Normalizing Feature Data

In this research, all feature values are normalized to the range of 0 and 1 so that it is easier for the next step of training machine learning models. For the features with a value larger than 1 or less than 0, we use the maximum possible value and minimum possible value in the training data set to normalize it.

#### 2.1.8 Data for training

We prepared the training dataset from the CASP dataset (CASP10-CASP12). The dataset is available at: http://predictioncenter.org/download_area/. Few protein decoys are filtered because of their low quality or failure of generating some of the features, and finally we have got 40,651 protein decoys for our machine learning model training process, which is described in the following section.

### 2.2 Machine learning model training

#### 2.2.1 Hierarchical Architecture

After all features are generated as described in the previous sections, the next step is to train machine-learning models. The traditional machine learning techniques have difficulty to train very deep networks, so deep learning techniques recently have been widely used in the bioinformatics field [22, 3, 11, 31]. We rely on the deep learning technique to understand the features and learn the pattern in the data, but it may not work when the number of input features is insufficient and may make incorrect predictions with small changes in the input feature [32]. We proposed a hierarchical architecture for machine learning model training, which is described in Figure 1. In order to make our machine learning model more robust, we select each pair of features from all input features and train a machine learning classifier on those two features to generate a higher level feature. By doing that, we will generate more features and could help to improve the performance of traditional machine learning model and deep learning technique.

**Figure 1:**
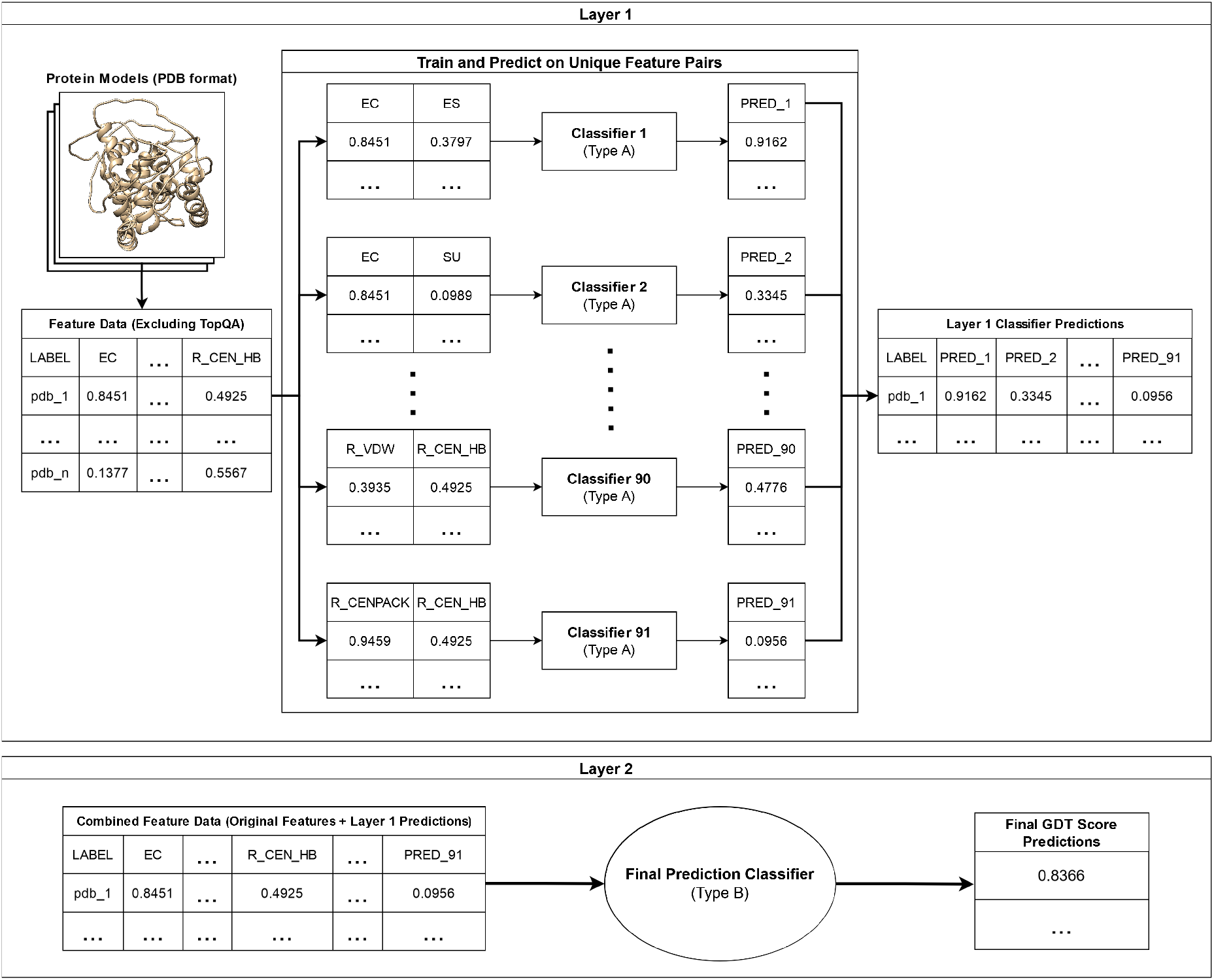
Overview of the hierarchical architecture of our machine learning model in training. Features are extracted from decoy models by our tool and exported to a CSV-like format. Every pair of features in the raw feature sets is then used to train separate machine learning classifiers of the same type and configuration (either Linear or Support Vector Regression in our experiments; see subsequent figures) before the next layer which will combine predictions made with these classifiers with the raw set of features to train the final machine learning classifier. Values in the figure are randomly generated for the sake of illustration.

First, several models of the same classifier type (“Type A”, see Figure 1) are trained on every possible pair of features in our data (without any redundancy; e.g. a model on feature 1 and feature 2, an additional model on feature 1 and feature 3, etc…), excluding those features generated from TopQA [27]. In total, 14 features are used to train the first layer of the hierarchical architecture. Every two of them are selected to train the type A model against the true GDT score of the decoy in the training data. The output of the first layer will be used for the second layer. This step can be repeated. There are 91 type A models to be trained in the first layer (as we are training on every possible combination of 2 features out of a pool of 14, see equation 1), so 91 additional features are generated from our raw features.

Second, we use the prediction generated in the previous step as part of the input features and train the final machine learning model of type “B” as the second layer of our hierarchical architecture. We only use two layers to demonstrate the effectiveness of our new architecture, but it can be expanded to more layers. The second layer will take the raw 14 features, 512 features from TopQA, and the 91 new generated features from the first layer of our hierarchical architecture as input. This makes 617 features in total for training of the second layer. We also experiment with just 105 features by excluding the ones from TopQA. For this paper, we explored Linear Regression (LR) and Support Vector Regression (SVR). SVR generated better results compared to Linear Regression (see discussion section), so we used SVR as the machine learning classifier for both architecture layers. The predicted output will be the final output of our new method. All 91 machine learning models from the first layer and one machine learning model from the second layer are stored for making predictions, where each prediction must fall between a range of 0 and 1. For instances where a prediction is above 1, the value is set to 0.9999, while predictions that are below 0 (negative values) get automatically set to 0.0001 to keep the predictions between our given range without losing the meaning of their original value.

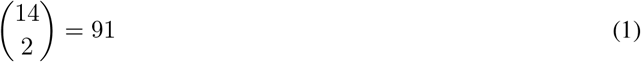

#### 2.2.2 Loss Function for Support Vector Regression

There are a lot of parameters for training the Support Vector Regression. The hyper-parameters (e.g., gamma, epsilon, etc.) of our Support Vector Regressions are optimized by the following two loss functions using GridSearchCV in the Sklearn tool (pedregosa2011scikit). The loss, which is the GDT score difference of the predicted “best” decoy and real best decoy in the input, is used as the metric for training:

- The difference between the true GDT of the true highest-ranked protein model and the true GDT of the predicted highest-ranked protein model (dubbed “Top-1 loss”).
- The difference between the average GDT of the true top 5 highest-ranking models and the average GDT of the predicted top 5 highest-ranking models (dubbed “Average loss”).

In both cases, a smaller loss value indicates better performance. The first one will focus on performance of the top 1 predicted model, and the second will focus on the performance of the top 5 predicted best model. The loss values shown in Figure 2 and Figure 3 are calculated as described in the second bullet-point, but as the difference in sums rather than the difference in averages.

**Figure 2:**
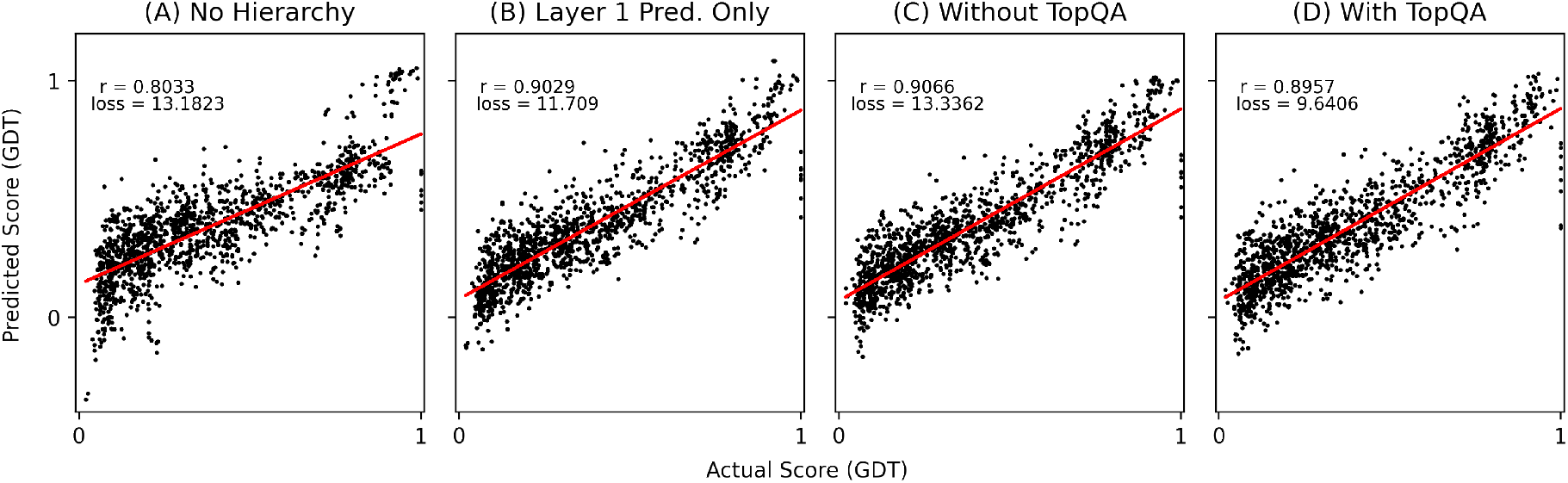
Correlations between actual GDT and predictions made by our method using Support Vector Regression as the type A classifier (see Figure 1), and Linear Regression as the type B classifier. **(A)** shows the performance of the final classifier trained on the original 14 features with no hierarchy architecture. **(B)** shows the performance of the final classifier using only layer 1 classifier predictions as input (so in total 91 features, see Figure 1). **(C)** demonstrates the performance of the final classifier using a combination of layer 1 classifier predictions and the original 14 features as input in layer 2, as shown in Figure 1 (so in total: 91 + 14 features). And finally, **(D)** shows the performance of the final classifier using the combination detailed in (C) with the addition of features generated through TopQA as input in layer 2 (a total of 91 + 14 + 512 features).

**Figure 3:**
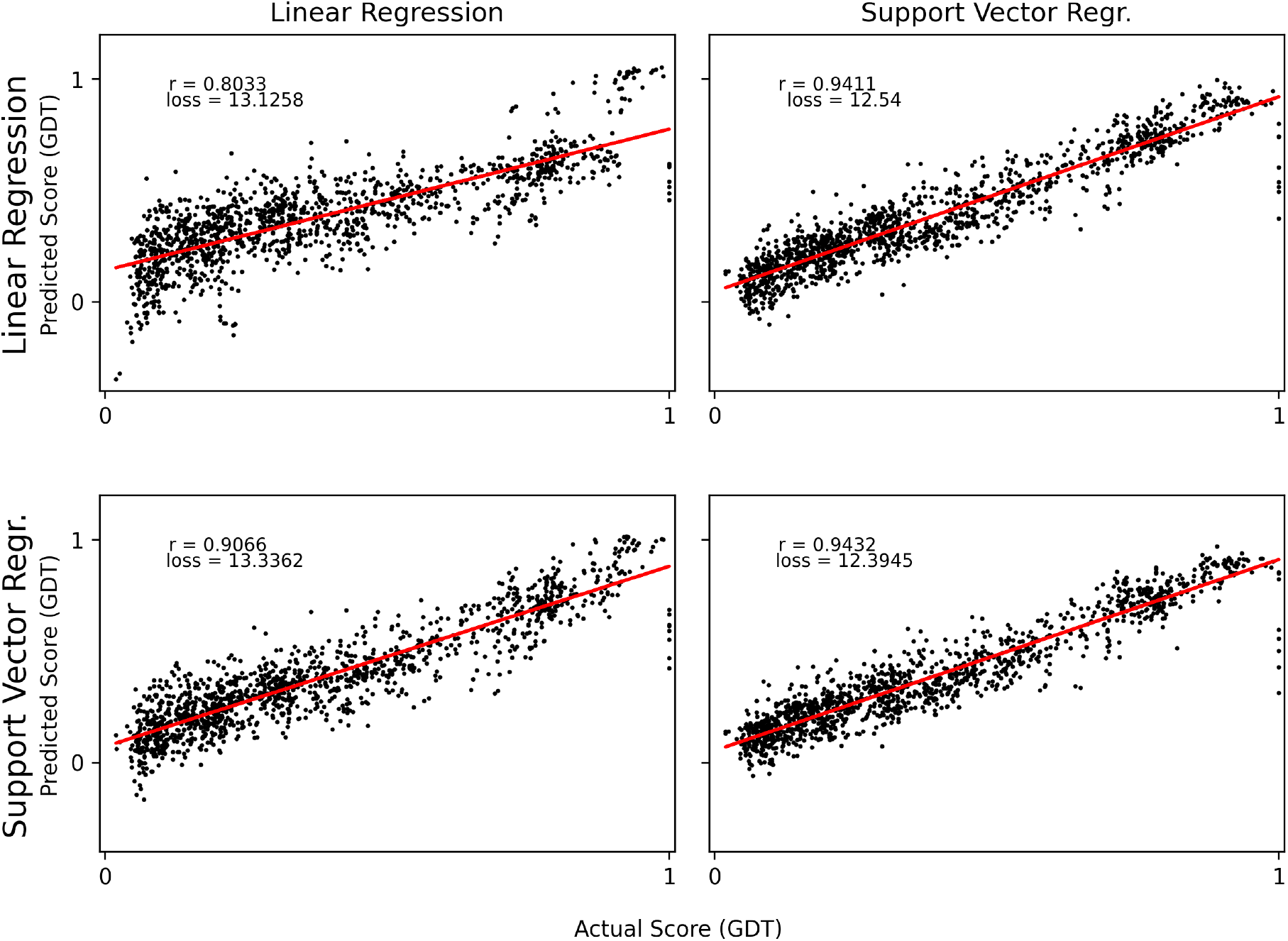
Correlations between actual GDT and predictions made by our method. Row labels represent layer 1 (Type A) classifiers, while column labels represent the final classifier (Type B) in layer 2. The architecture here uses the same set of features as plot (C) in Figure 2.

To rank protein decoys, the final version of our SynthQA tool uses a combination of SVR models optimized with the two loss functions listed above. The tool first ranks models by the average loss method, and then generates a seperate ranking list using the top-1 loss method to identify the highest-scoring decoy. Then, our method will move that “top-1” decoy to the top of the average loss rankings by

## 3 Discussion

In order to validate the effectiveness of our new hierarchical architecture, we evaluate the performance of Linear Regression (LR) and Support Vector Regression (SVR) classifiers with and without our new architecture. Figure 2 shows the correlation of actual GDT and predictions made by our tool on structures from the CASP 12 dataset (4,713 decoys) with different configurations. As we can see in Figure 2, Figure 2(A) possesses the lowest Pearson Correlation of 0.8033 with a loss value of 13.1823. Figure 2(B) demonstrates that the addition of our hierarchy increases this correlation to 0.9029, and decreases the loss to 11.709. We also see that in Figure 2(C), the addition of the original features provided at the beginning of our architecture combined with predictions made in layer 1 as input for the final classifier in layer 2 continues to increase the correlation to 0.9066 at the cost of increasing loss to 13.3362. We also tried combining the features used in Figure 2(C) with features generated with TopQA to use as input for the final classifier in layer 2, and found that the correlation of predictions and actual GDT is 0.8957 with a loss value of 9.6406, shown in Figure 2(D). This proves the effectiveness of our hierarchical architecture to improve the performance of traditional machine learning techniques by generating more features and training them layer by layer. In our experiments, we have found that the hierarchical architecture didn’t work well to improve the LR because our architecture connected new generated features in a linear way which didn’t influence the linear regression by much.

In the final version of our method, we have chosen to prioritize higher correlation over minimized loss. This would mean that (C) in Figure 2 would be our optimal configuration of choice. The results in Figure 2(D) have shown that using TopQA did not improve the correlation, and so the TopQA features were ultimately not used.

We did another experiment that compares the performance of our method with different type A and type B classifiers (see Figure 1) to see which combination would work best between LR and SVR classifiers. Using the (C) configuration from Figure 2, the plots in Figure 3 compare the hierarchy’s performance with the following type A and type B classifiers: LR to LR, LR to SVR, SVR to LR, and SVR to SVR. We can see in Figure 3 that the highest correlation of 0.9432 is obtained by using Support Vector Regression as both the type A and type B classifier in both layers of the architecture. This configuration also yields the smallest loss out of the four shown, with a value of 12.3945. This lines up with our understanding of Support Vector Regression being more suitable and accurate for data containing a large amount of features.

In summary, SVR performs better than LR in our experiment. So it is recommended to use SVR in both layers of our hierarchical architecture. The hierarchical architecture is effective at generating additional features and improving the performance of traditional machine learning methods without needing extra base features. Moreover, our method also shows that using the relationship between features as proposed in our hierarchical architecture is helpful and we can add more layers in the hierarchical architecture to further improve the performance.

### Benchmark on CASP 14

In order to benchmark the performance of our new tool SynthQA, we tested SynthQA on the latest CASP 14 data sets and compared with a few top-performing single model QA methods. We use the standard evaluation metrics: per-target average Pearson correlation, per-target Kendall Tau correlation, per-target average Spearman’s correlation, per-target average loss, overall Pearson correlation, overall Kendall Tau correlation, and overall Spearman’s correlation. Figure 4 illustrates the performance of our method and several other methods on Stage 1 CASP 14 data sets. As we can see in the figure, SynthQA is the second best on the overall Pearson correlation, Kendall tau correlation, and Spearman’s correlation, compared to other top-performing methods like DeepQA, VoroMQA, and SMOQ. If we evaluate the performance based on the per-target metric, our method is not performing well based on the correlations like per-target average Pearson correlation, per-target Kendall Tau correlation, or per-target average Spearman’s correlation. This could because we use the loss as the metric to train our method instead of the correlation. However, the loss of our method is one of the best (0.09, and the other method VoroMQA also has the loss 0.09), which shows that the quality of top 1 model selected by our method is very good. We also evaluated the performance based on LDDT score and didn’t notice a big change for the performance of our method. In addition, we evaluated the performance on Stage 2 CASP 14 datasets which is shown in Figure 5. Our method did a better job on the per-target metric, we are slightly better than SMOQ on the per-target Kendall Tau correlation and per-target average Spearman’s correlation. The per-target loss of our method on Stage 2 is 0.14, better than DeepQA and SMOQ, and the same as the best method VoroMQA using GDT as the metric. When we use LDDT score as the metric for our evaluation, VoroMQA has 0.08 average per-target loss, a little bit better than us with a 0.09 per-target loss. Overall, our experiment shows that our method is comparable with other top performing methods and adept at selecting the top 1 model from the protein decoys model pool.

**Figure 4:**
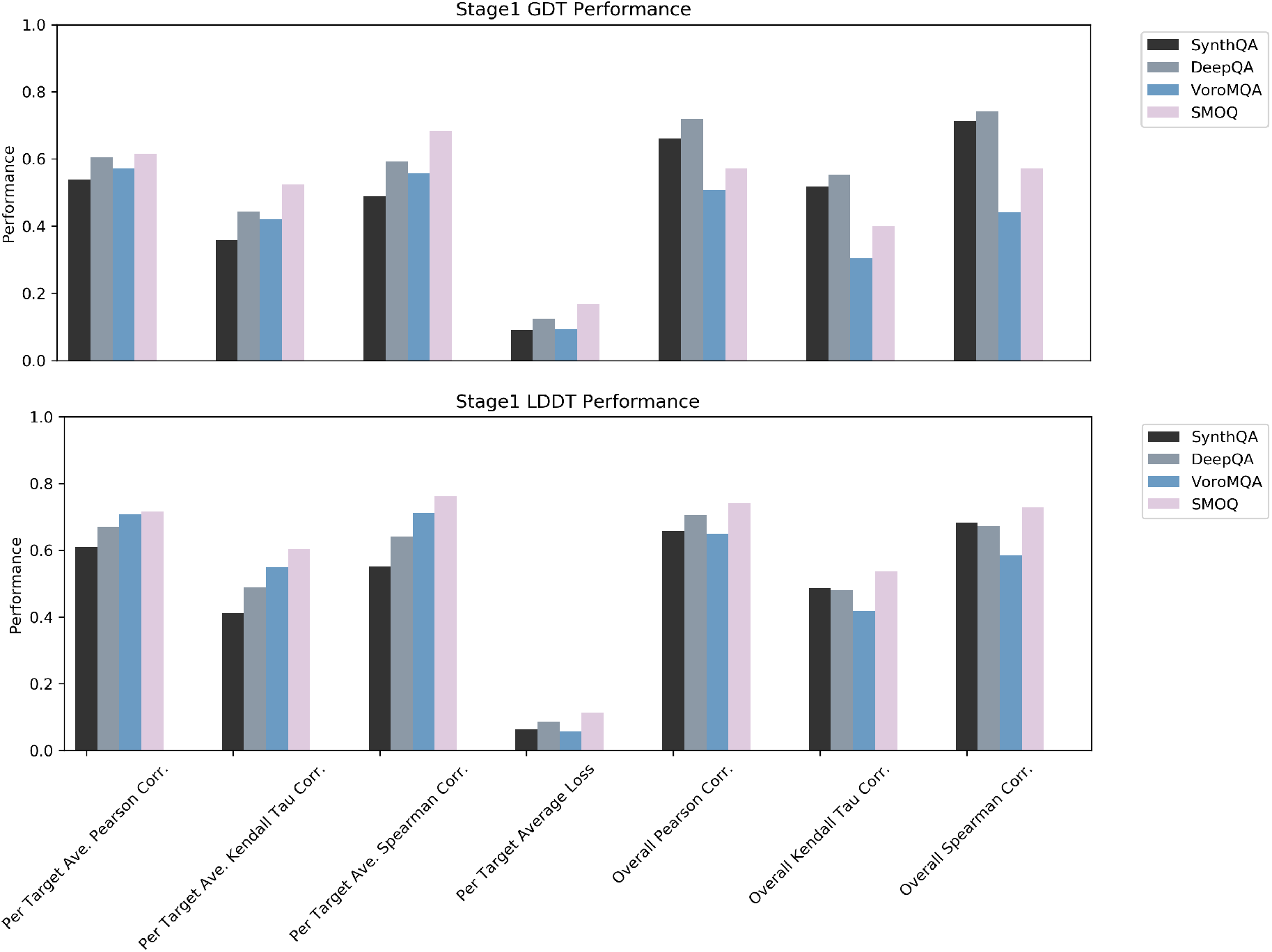
Performance of our method and several other top-performing method on Stage 1 (sel20) CASP 14 datasets using GDT and LDDT respectively. The evaluation metrics we used is per-target average Pearson correlation, per-target Kendall tau correlation, per-target average Spearman’s correlation, per-target average loss, overall Pearson correlation, overall Kendall tau correlation, and overall Spearman’s correlation. For the loss, the performance is better when it is smaller, and for the correlation, the larger the better.

**Figure 5:**
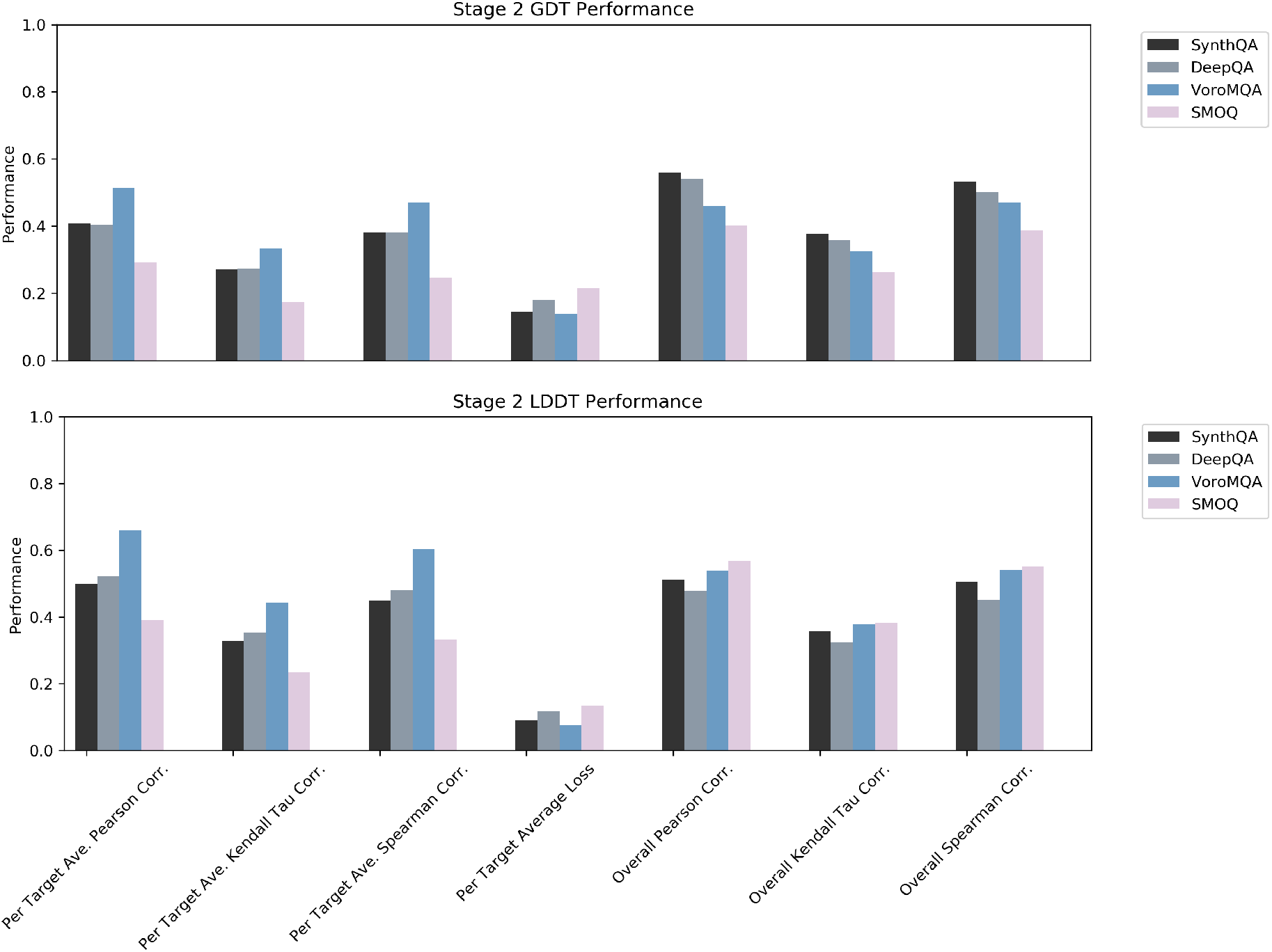
Performance of our method and several other top-performing method on Stage 2 (top150) CASP 14 datasets using GDT and LDDT respectively. The evaluation metrics we used is per-target average Pearson correlation, per-target Kendall tau correlation, per-target average Spearman’s correlation, per-target average loss, overall Pearson correlation, overall Kendall tau correlation, and overall Spearman’s correlation

#### 3.0.1 Performance of features used in SynthQA

It is interesting to find out the performance of each feature used in SynthQA and compare it with SynthQA. We used CASP 14 targets and benchmark the performance of each feature used in SynthQA. As we can see in Figure 6, SynthQA consistently outperforms each of the 14 features in CASP 14 datasets, and especially the loss of SynthQA is much better than each individual single feature, which shows the effectiveness of our new hierarchical architecture.

**Figure 6:**
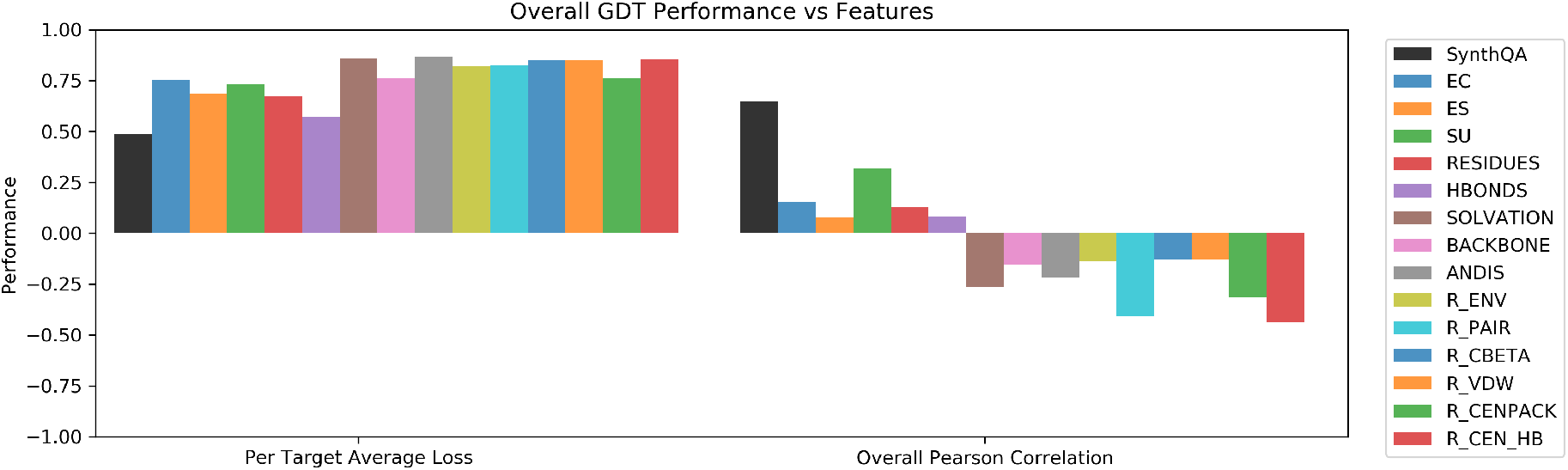
Overall performance of all 14 features and SynthQA on all models in CASP 14 datasets. GDT score is used as the metric for the calculation.

## Conclusion

In this paper, we propose a new protein model quality assessment tool, SynthQA, which uses multi-scale features and a new hierarchical architecture that can generate additional features for training machine learning techniques. Our experiment shows that SynthQA is comparable to other state-of-the-art methods for ranking and selecting protein structure decoys and can be a useful tool for protein structure prediction. In the future, we plan to add more layers in our hierarchical architecture and add more features from the protein sequence. We also hope to experiment with different machine learning techniques to use in-place of Support Vector Regression, as well as explore the possibility of deep-learning integration.

## Acknowledgements

This work is supported by the Natural Sciences Undergraduate Research Program at Pacific Lutheran University to R.C‥ This material is based upon work is also supported by the National Science Foundation under Grant No. 2030381 and the graduate research award of Computing and Software Systems division at University of Washington Bothell to D.S‥ Any opinions, findings, and conclusions or recommendations expressed in this material are those of the authors and do not necessarily reflect the views of the National Science Foundation.

